# Multidimensional super-resolution microscopy unveils nanoscale surface aggregates in the aging of FUS condensates

**DOI:** 10.1101/2023.07.12.548239

**Authors:** Changdong He, Chun Ying Wu, Wan Li, Ke Xu

## Abstract

The intracellular liquid-liquid phase separation (LLPS) of biomolecules gives rise to condensates that act as membrane-less organelles with vital functions. FUS, an RNA-binding protein, natively forms condensates through LLPS and further provides a model system for the often disease-linked liquid-to-solid transition of biomolecular condensates during aging. However, the mechanism of such maturation processes, as well as the structural and physical properties of the system, remain unclear, partly attributable to difficulties in resolving the internal structures of the micrometer-sized condensates with diffraction-limited optical microscopy. Harnessing a set of multidimensional super-resolution microscopy tools that uniquely map out local physicochemical parameters through single-molecule spectroscopy, here we uncover nanoscale heterogeneities in the aging process of FUS condensates. Through spectrally resolved single-molecule localization microscopy (SR-SMLM) with a solvatochromic dye, we unveil distinct hydrophobic nanodomains at the condensate surface. Through SMLM with a fluorogenic amyloid probe, we identify these nanodomains as amyloid aggregates. Through single-molecule displacement/diffusivity mapping (SM*d*M), we show that such nanoaggregates drastically impede local diffusion. Notably, upon aging or mechanical shears, these nanoaggregates progressively expand on the condensate surface, thus leading to a growing low-diffusivity shell while leaving the condensate interior diffusion-permitting. Together, beyond uncovering fascinating nanoscale structural arrangements and aging mechanisms in the single-component FUS condensates, the demonstrated synergy of multidimensional super-resolution approaches in this study opens new paths for understanding LLPS systems.

## Text

The past decade has witnessed the fast growth of research on microdroplet-like “condensates” generated and maintained *via* the liquid-liquid phase separation (LLPS) of biomolecules^1-4^. By bringing together relevant molecular players, including proteins and nucleic acids, and modulating local physicochemical parameters, *e*.*g*., diffusivity and hydrophobicity, such “membrane-less organelles” give rise to vital cellular functions^5^, *e*.*g*., signaling^6,7^ and gene control^8-10^.

The RNA-binding protein FUS (fused in sarcoma) and related proteins serve as model systems for LLPS and biomolecular condensates^11-19^. Formed through LLPS owing to intrinsically disordered low-complexity domains, FUS condensates are further known for their gradual transition from an initially liquid state to a solid or glassy state through aging^11,14,16,19^, a process accelerated by mutations linked to neurodegenerative diseases^11,12,15,18^. Related aging/maturation processes have been reported in numerous LLPS systems, including other neurodegenerative-disease players such as hnRNPA1^20^, Tau^21,22^, TDP-43^23^, and α-synuclein^24^. However, the mechanism of such processes, as well as the overall structural and physical properties of the system, remain unclear. For example, for single-component LLPS condensates, whereas it is common to assume that each microdroplet adopts a homogenous phase, recent theoretical and experimental work points to potential intra-condensate inhomogeneity, *e*.*g*., the gradual transformation of FUS LLPS condensates to liquid-core/gel-shell multiphase architectures through aging^19,25^.

The detection and elucidation of possible nanoscale heterogeneities in the often micrometer-sized condensate droplets are impeded by the ∼300 nm resolution of diffraction-limited optical microscopy. The past decade has seen substantial advances in super-resolution microscopy (SRM), including single-molecule localization microscopy (SMLM), which, by super-localizing the positions of millions of individual molecules over consecutive camera frames, routinely achieves ∼20 nm spatial resolution^26-29^. However, limited SRM applications have been demonstrated for LLPS condensates^30-32^. Moreover, intra-condensate heterogeneities may present as varied local physicochemical parameters independent of morphology or molecular distributions, which may not be resolvable by enhancing the spatial resolution alone, as will be demonstrated in our data below.

We recently developed an arsenal of *functional* SRM methods that map out physicochemical parameters at the nanoscale by tapping into the high-dimensional information space of single-molecule fluorescence^29,33-36^. Spectrally resolved SMLM (SR-SMLM) detects local parameters through the emission spectra of single probe molecules^33-35^. Single-molecule displacement/diffusivity mapping (SM*d*M) maps out diffusivity with high spatial resolutions to unveil local molecular states and interactions^36-39^. Together, these approaches have provided valuable insights into diverse cellular^34,36-39^ and *in vitro*^40-42^ systems.

Here we harness these technical advances to uncover nanoscale heterogeneities in the aging process of LLPS condensates, and we focus on probing the local chemical polarity, protein states, and diffusivity for the FUS system. We thus unexpectedly unveil distinct nanodomains of low chemical polarity at the condensate surfaces, and further show that these nanodomains are due to amyloid fibril aggregates and drastically impede local diffusion. Moreover, we find that such nanoaggregates progressively expand on the condensate surface upon aging or mechanical shears, thus leading to a growing low-diffusivity shell while leaving the condensate interior diffusion-permitting.

## Results and discussion

SR-SMLM achieves high-throughput single-molecule spectroscopy by integrating SMLM with a wide-field dispersion scheme (Fig. 1a)^35^. The resultant spectrally resolved SRM images allow functional encoding with environment-sensing probes, *e*.*g*., Nile Red for probing local chemical polarity^34,43^. The solvatochromic dye Nile Red is nonfluorescent in the aqueous phase but exhibits fluorescence switch-on and spectral blueshift in hydrophobic (low chemical polarity) environments, and so has been valuable for the (SR-)SMLM of lipid membranes^34,43,44^ and protein aggregates^43,45^.

**Figure 1.**
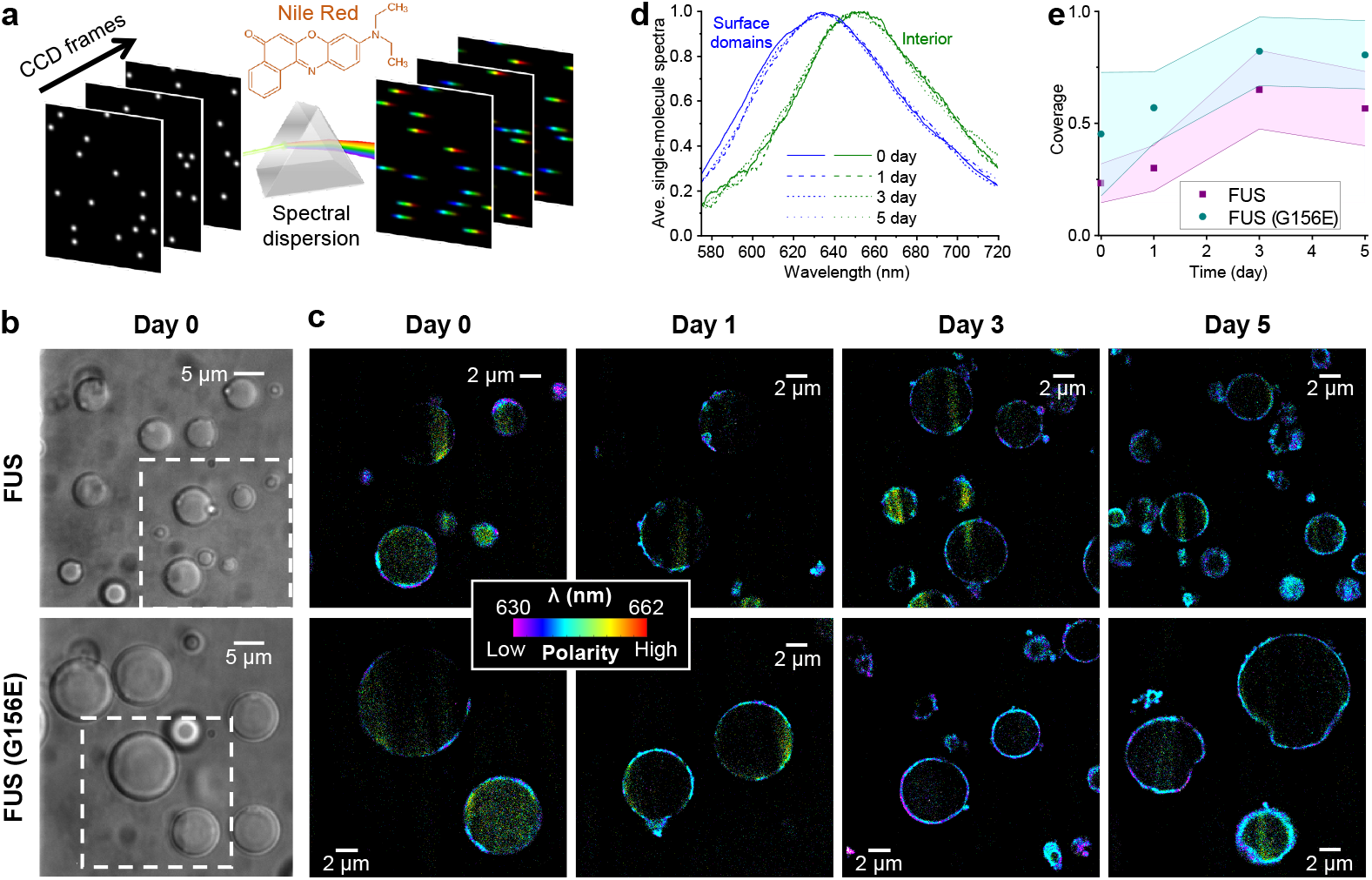
SR-SMLM with Nile Red unveils growing hydrophobic (low chemical polarity) nanodomains at the FUS condensate surface. **a**. Schematics: Single-molecule fluorescence is spectrally dispersed in the wide field, and the resultant single-molecule spectra are accumulated over many camera frames to generate SR-SMLM images. **b**. Brightfield micrographs of condensates formed by wild-type (top) and G156E-mutant (bottom) FUS. **c**. Nile Red SR-SMLM images for FUS (top) and FUS(G156E) (bottom) condensates aged for different days. Color presents the mean wavelength of the local single-molecule spectra (color scale bar), which reflects the local chemical polarity. The Day 0 images correspond to the boxed regions in (b). **d**. Averaged single-molecule spectra at the condensate surface hydrophobic domains (blue) and the condensate interior (green), for FUS condensates aged for different days (different line shapes). **e**. Time-dependent surface coverage of the Nile Red-visualized hydrophobic domains over aging. Data points and shades present the averages and standard deviations of individual condensate droplets (∼10 for each data point), respectively.

We employed Nile Red-based SR-SMLM to interrogate local chemical polarity in FUS condensates. To this end, we prepared wild-type and mutant FUS proteins and induced condensates following typical protocols^11,46^. Micrometer-sized droplets were formed as expected (Fig. 1b). As we illuminated a 561-nm laser a few micrometers into the sample and kept the focal plane at that depth to image a cross-section of the condensate droplets, stochastic local bursts of single-molecule fluorescence were observed, corresponding to individual Nile Red molecules transiently encountering a hydrophobic (low chemical polarity) phase in the sample. The wide-field fluorescence signal was split into two optical paths, with one spectrally dispersed, so single-molecule images and spectra were concurrently recorded for different molecules in the view^34,35^. Accumulating the single-molecule images and spectra over ∼10^4^ frames generated spectrally resolved super-resolution images (Fig. 1c), in which color presented the spectral mean of locally accumulated molecules^34,35^, hence the local chemical polarity.

We thus unveiled intriguing nanostructures in the condensates. Notably, the detected Nile Red signal highlighted discrete segments ∼100 nm in apparent thickness at the condensate surface (Fig. 1c and Fig. S1). Averaging the local single-molecule spectra at these segments showed a substantial blueshift to an ∼635 nm peak when compared to the ∼650 nm peak averaged from the sparsely detected single-molecule signal at the condensate interior (Fig. 1d and Fig. S1). The latter value matches that found in highly hydrated environments, whereas the former blue-shifted result is consistent with the more hydrophobic environments seen at protein aggregates^47,48^.

Repeating Nile Red SR-SMLM over 5 days next showed that whereas the hydrophobic domains initially occurred as sporadic segments, covering ∼23% of the wild-type FUS condensate surface, the surface coverage increased gradually, reaching ∼60% after 3 and 5 days (Fig. 1ce and Fig. S1). The apparent thickness of this layer remained ∼100 nm over the process, suggesting that the hydrophobic domains expanded laterally on the condensate surface with a fixed thickness.

We compared results with the FUS(G156E) mutant, an ALS/FTD-associated mutation known to accelerate the condensate solidification process^11,15^. We thus observed similar hydrophobic segments ∼100 nm in thickness at the condensate surfaces (Fig. 1c and Fig. S2), yet noting a higher surface coverage of ∼45% in the as-prepared condensates, and finding this coverage grew to ∼80% by Day 3 and Day 5 (Fig. 1ce). The latter higher surface coverages further coincided with when the condensates started to lose their rounded shapes to become irregular in morphology. The averaged Nile Red single-molecule spectra at the surface segments and the condensate interior (Fig. S2) were both indistinguishable from that of wild-type FUS, suggesting similar protein states.

Thus, SR-SMLM unveiled the gradual lateral expansion of ∼100 nm thick hydrophobic nanodomains at the surface of FUS condensates, and this process was accelerated by the G156E mutation. Meanwhile, the condensate interior remained highly hydrated with high chemical polarity throughout the aging process.

For *in vitro* protein systems, Nile Red fluorescence switch-on and spectral blueshifts often occur at amyloid fibrils^43,47,48^. To examine this possibility, we utilized CRANAD-2 (Fig. 2a), a far-red fluorogenic probe with high affinity and specificity to amyloid fibrils^49,50^. A recent study has demonstrated CRANAD-2 for the SMLM of α-synuclein fibrils^50^, in which the stochastic amyloid binding of individual CRANAD-2 molecules gives bursts of single-molecule fluorescence.

**Figure 2.**
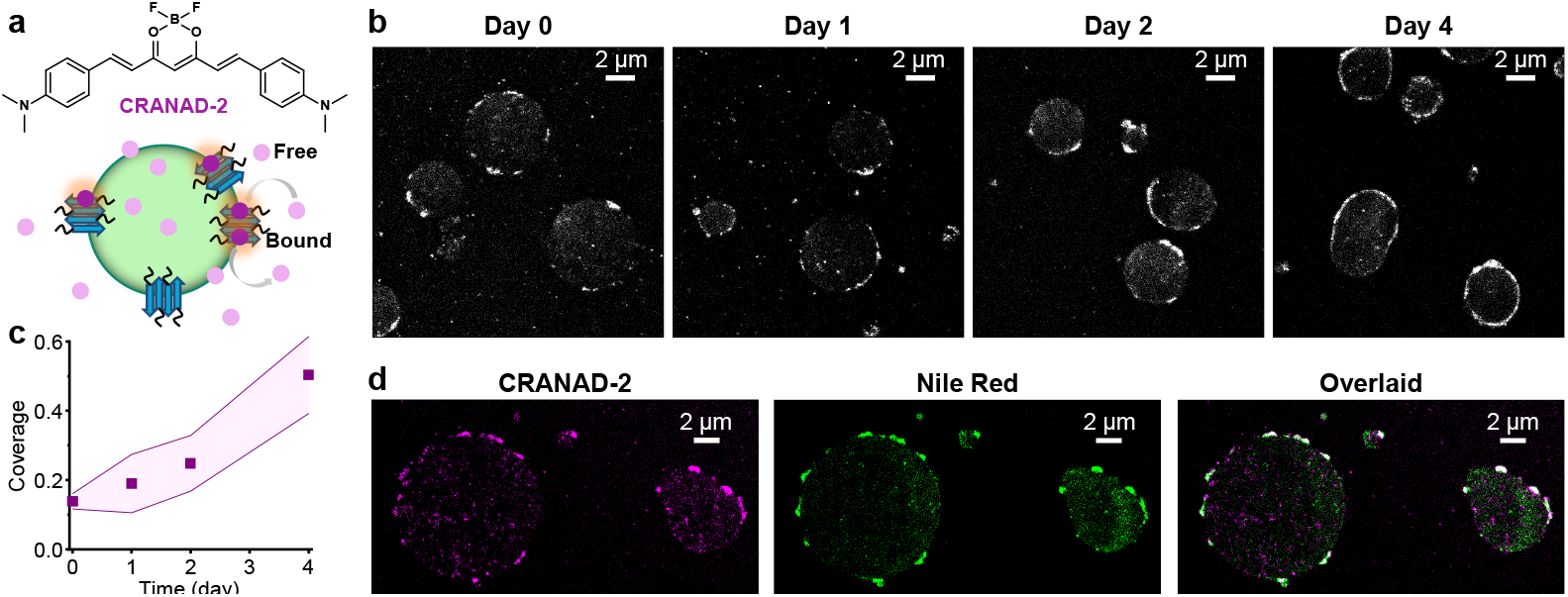
SMLM with CRANAD-2 indicates the gradual formation of amyloid fibril nanoaggregates at the FUS condensate surface. **a**. CRANAD-2 chemical structure and mode of action: CRANAD-2 is nonfluorescent in the aqueous phase but turns on fluorescence upon binding to amyloid fibrils in the condensates. **b**. Representative SMLM images of CRANAD-2, for FUS condensates aged for different days. **c**. Time-dependent surface coverage of CRANAD-2 SMLM signal at FUS condensates over aging. Data points and shades present the averages and standard deviations of individual condensates (∼10 for each data point), respectively. **d**. Two-color SMLM for a FUS LLPS sample loaded with both CRANAD-2 and Nile Red, shown as separate (CRANAD-2: magenta, Nile Red: green) and overlaid (colocalization: white) images.

Notably, as we applied CRANAD-2 to the FUS condensate system and performed SMLM under 642 nm excitation, we identified surface nanostructures consistent with the hydrophobic nanodomains observed above with Nile Red-based SR-SMLM. Discrete segments ∼100 nm in thickness were noted at the condensate surfaces (Fig. 2b), and they similarly increased in surface coverage while maintaining their thinness when aged over days (Fig. 2bc).

As CRANAD-2 and Nile Red are spectrally distinct, we further loaded a sample with both dyes and sequentially performed SMLM for the two using their respective fluorescence filter cubes and excitation lasers. We thus found that the CRANAD-2 and Nile Red SMLM results exhibited good colocalization, both highlighting the same surface nanodomains (Fig. 2d).

Thus, with two dyes that respectively probe local chemical polarity and protein states, our SRM results indicate the gradual lateral expansion of a thin hydrophobic shell in the aging of FUS condensates through the formation of amyloid fibril aggregates.

We next inquired whether the surface aggregates we uncovered could alter local physical environments and hence the behavior of guest molecules. Here we focus on mapping molecular diffusivity, which both reflects and modulates intermolecular interactions. For this task, we turn to Cy3B (Fig. 3b), a consistently bright dye often used in single-molecule microscopy^51,52^, contrasting with Nile Red and CRANAD-2 above only highlighting hydrophobic or amyloid phases. With a molecular weight of 560 Da, Cy3B serves as a good proxy for the mobility of similarly sized intracellular signaling molecules. Through SM*d*M, we have recently quantified the fast diffusion of this dye in water, hydrogels, and mammalian cells^39,42^.

**Figure 3.**
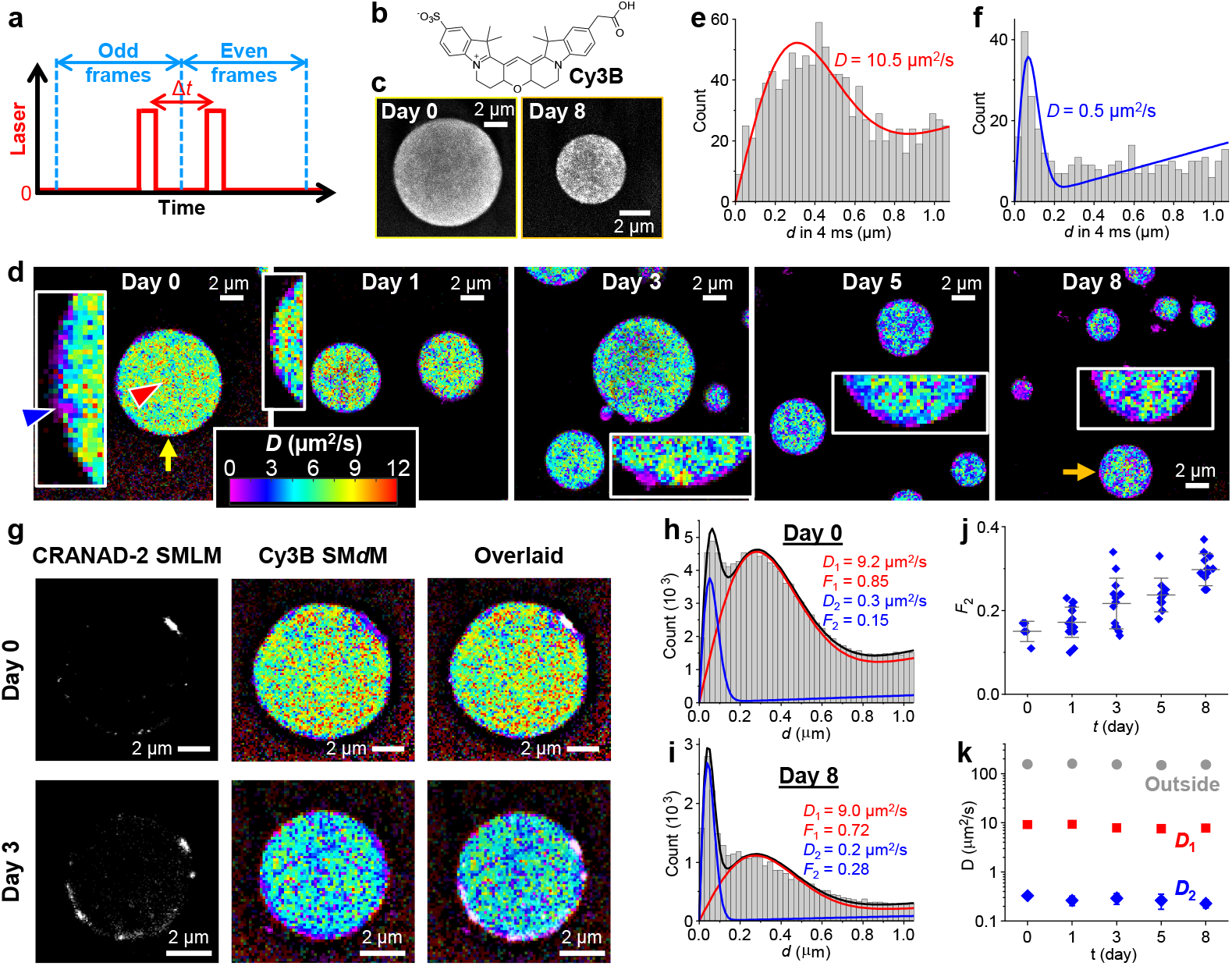
SM*d*M with Cy3B shows impeded diffusion in the FUS condensates and further large drops in diffusivity at the surface aggregates. **a**. Scheme of SM*d*M. Paired stroboscopic excitation pulses are repeatedly applied in tandem across odd-even camera frames, so that single-molecule displacements are captured for the time window defined by the separation between the tandem pulses Δ*t* rather than the camera framerate. **b**. Chemical structure of Cy3B. **c**. SMLM images based on the localized positions of single Cy3B molecules, for FUS condensates as prepared (left) and aged for 8 days (right). **d**. Color-coded SM*d*M super-resolution *D* maps of Cy3B in FUS condensates aged for different days, obtained by spatially binning the accumulated single-molecule displacements onto a 120 nm grid, and then fitting the distribution in each bin to extract the local *D* for color rendering. The condensates indicated by the yellow and orange arrows correspond to the ones shown in (c). Insets: zoom-ins of the condensate surfaces, highlighting local diffusion slowdowns. **e**,**f**. Histograms: Local distributions of the accumulated single-molecule displacements at Δ*t* = 4 ms, for two regions pointed to by the red and blue arrowheads in (d), respectively. Red and blue curves: Fits to our diffusion model, with resultant *D* values marked in the plots. **g**. Sequentially acquired CRANAD-2 SMLM images and Cy3B SM*d*M *D* maps for FUS condensates as-prepared (top) and aged 3 days (bottom), shown as separated and overlaid images. **h**,**i**. Two-component fits (red curve: fast component; blue curve: slow component; black curve: sum) to the single-molecule displacements (histograms) collected at the FUS-condensate interior at Day 0 (top) and Day 8 (bottom). Resultant *D* values and fractions of the two components are marked in the plots. **j**. Fraction of the low-diffusivity component, as obtained above, for the interior of FUS condensates aged for different days. Each data point corresponds to one individual condensate. **k**. Fitted *D* values for the fast (red) and slow (blue) components, as obtained above, for the interior of FUS condensates aged for different days, versus *D* values in the dilute phase outside the condensates (gray). Error bars in (j,k): standard deviations between individual condensates.

We thus similarly prepared FUS condensates as above and added Cy3B. For SM*d*M, paired stroboscopic excitation pulses were repeatedly applied in tandem across odd-even camera frames (Fig. 3a), so that single-molecule images due to Cy3B molecules stochastically diffusing into the focal plane were captured, and their displacements were detected over the time window defined by the pulse separation^36^.

SMLM based on the super-localized single-molecule positions over ∼10^4^ paired frames showed that Cy3B was concentrated in the droplet-like condensates, where it distributed relatively homogeneously at the nanoscale (Fig. 3c). Cy3B is moderately hydrophilic; its enhanced presence in the condensates over the dilute phase may be related to its substantially reduced diffusivity in the condensates (below).

Spatially binning the accumulated single-molecule displacements with a 120 nm grid next enabled^29,36^ fitting the distribution of displacements in each spatial bin to extract the local diffusion coefficient *D*. The resultant color-coded super-resolution *D* maps (Fig. 3d) and local distribution of single-molecule displacements (Fig. 3e for the region indicated by the red arrowhead in Fig. 3d) showed that the fastest diffusing regions inside the condensates reached *D* ∼10 μm^2^/s. Whereas such diffusivity would have been too fast to follow with traditional single-molecule tracking, it is only ∼3% of Cy3B in water (∼340 μm^2^/s)^39,42^. We have recently shown that in hydrogels, Cy3B only exhibits mild (∼15%) reductions in *D* at ∼6 wt% polymer contents^42^, and that in mammalian cells, Cy3B diffusion is only reduced by ∼25% owing to the modestly higher viscosity of the cytoplasm over water^39^. Thus, our results suggest high macromolecular crowding and/or effective viscosity for the 560 Da Cy3B in the FUS condensates. Such markedly reduced mobilities may be functionally significant for the condensates to act as concentrating hubs and interaction sites for biomolecules^2-5^.

A closer examination of the super-resolution *D* maps noted nanoscale segments of even greater drops in diffusivity at the condensate surface. This observation is challenging: As a 120 nm grid was implemented to warrant enough single-molecule displacements in each spatial bin for fitting, the ∼100 nm thick shell we identified above at the condensate surface would be pixelated into just one or two bins in thickness. Nonetheless, facilitated by the occasionally observed nanoscale inclusions, regions with drastically reduced *D* of ∼0.5 μm^2^/s were identified (blue arrowhead in Fig. 3d inset; Fig. 3f). Such extremely low *D* at the condensate surface likely corresponds to Cy3B bound to the solid-like aggregates we identified above. Indeed, as we monitored the FUS condensates over days with Cy3B SM*d*M, we observed that the low-diffusivity domains gradually increased their coverage at the surface (Fig. 3d and insets), although quantification was difficult due to the pixelation effects.

To directly compare with surface aggregates, we added both CRANAD-2 and Cy3B to new samples, and sequentially performed SMLM and SM*d*M for the two dyes using their respective fluorescence filter cubes and excitation lasers. We thus found that the SM*d*M-resolved Cy3B slowdown regions at the condensate surface correlated well with the CRANAD-2 SMLM signal (Fig. 3g). Thus, the surface amyloid fibril aggregates are characterized by drastically lowered local diffusivities.

Diffusion inhomogeneities were also noted inside the condensates. Interestingly, the collective distribution of single-molecule displacements from the condensate interior showed two peaks (Fig. 3h for Day 0). We thus introduce a two-component fitting model to extract two *D* values and their respective fractions (methods). Notably, the resultant *D*_1_ = 9.2 μm^2^/s and *D*_2_ = 0.3 μm^2^/s were respectively similar to the *D* values observed above for the fast-diffusion regions and surface aggregates. The fraction of slow diffusion was *F*_2_ ∼15% for the as-prepared condensates. However, the SM*d*M data may have overrepresented the slower population, which stays better in focus between the tandem excitation pulses.

Analyzing the SM*d*M data of the condensate interior over aging next showed that both *D*_1_ and *D*_2_ remained largely unvaried (Fig. 3ik), yet the apparent fraction of the slow population gradually increased from ∼15% to ∼30% in 8 days (Fig. 3ij). We also examined Cy3B diffusion in the dilute phase outside the condensates and found *D* ∼150 μm^2^/s, expected with the increased medium viscosity from dextran addition. This *D* value remained constant over 8 days (Fig. 3k), suggesting no substantial changes in the dilute-phase properties in aging.

Together, our SM*d*M results suggest that whereas the surface aggregates are mostly nonmobile, the condensate interior is dominated by a fluidic phase plus a small low-diffusivity fraction. The low-diffusivity fraction likely corresponds to nanoscale aggregates inside the condensates, which, while not readily resolved in SM*d*M with the 120 nm bin sizes convolved with the different depths within the focal range, stand out through single-molecule displacement distribution analysis. Over aging, the fluidic phase maintains its liquid nature with a constant diffusivity, yet the low-diffusivity fraction steadily increases its presence, suggesting the gradual enrichment of aggregates, which may further accumulate on the condensate surface to expand the shell.

A recent study^17^ has shown that mechanical shear promotes the liquid-to-solid transition of FUS condensates and results in enhanced fluorescence of Thioflavin T, a common amyloid probe. To test if this behavior could be related to our above-observed gradual formation of amyloid-like aggregates at the FUS condensate surface in aging, we prepared FUS LLPS samples in which we imposed mechanical shear by repeated pipetting.

For samples pipetted 20 times, we thus observed significantly increased coverage of amyloid nanoaggregates at the FUS condensate surface, as visualized by the colocalization of CRANAD-2 SMLM signal and reduced local diffusivity of Cy3B in the SM*d*M *D* map (Fig. 4a). Quantification indicated that the CRANAD-2 SMLM signal covered ∼60% of the condensate surface (Fig. 4c), substantially higher than that of the undisturbed samples (Fig. 2c) as prepared (∼20%) or aged for 4 days (∼50%). Moreover, irregular-shaped nanoaggregates were noted extruding out of the condensate surface (arrows in Fig. 4a).

**Figure 4.**
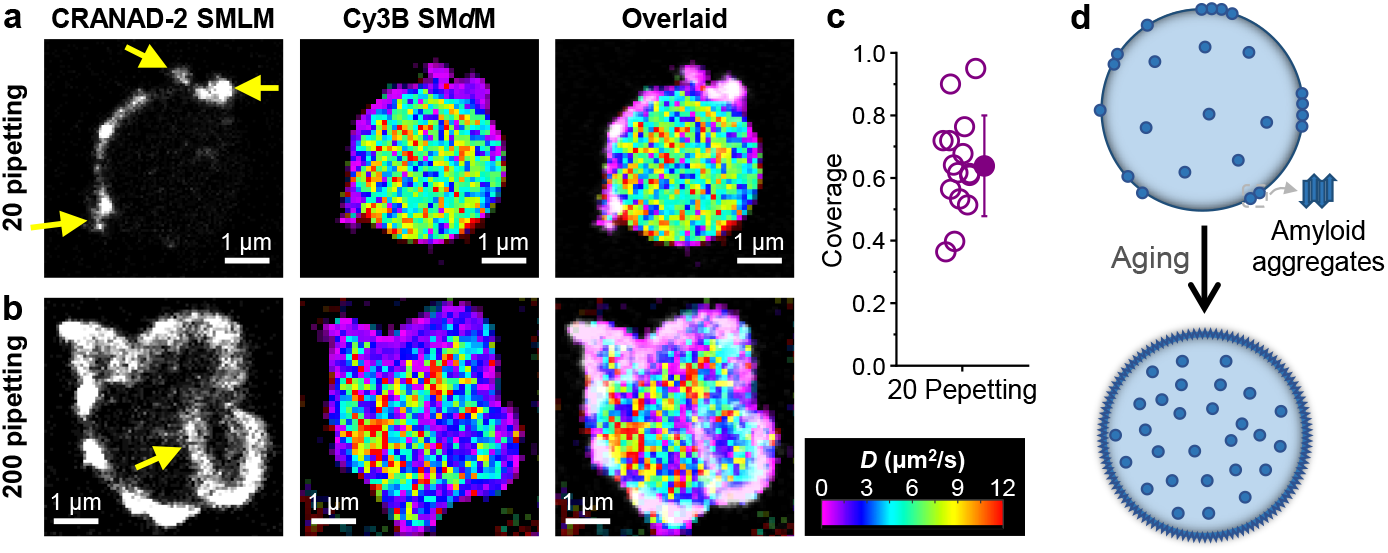
Mechanical shear induces similar amyloid fibril aggregates at the FUS condensate surface. **a**,**b**. Sequentially acquired CRANAD-2 SMLM images and Cy3B SM*d*M *D* maps, and overlaid images, for FUS condensates subjected to 20 (a) and 200 (b) times of pipetting. Arrows in (a,b) point to extrusions and intrusions of the surface aggregates. **c**. Surface coverage of CRANAD-2 SMLM signal for FUS condensates pipetted 20 times. Open circles: Individual condensates. Filled circle and error bar: Average and standard deviation. **d**. Model: Enrichment of amyloid aggregates in the condensates and accumulation at the surface during aging.

For samples pipetted 200 times, the condensates lost their droplet-like rounded shapes, but appeared as granules of irregular shapes. Interestingly, CRANAD-2 SMLM (Fig. 4b) showed high coverage of the granule surfaces to form a shell thickened to ∼400 nm, yet signal from the granule interior remained low, except for aggregate intrusions from the surface (arrow in Fig. 4b). Meanwhile,

Cy3B SM*d*M showed impeded diffusion for a thickened shell that corresponded well with the CRANAD-2 SMLM signal (Fig. 4b). Notably, diffusion at the granule interior remained reasonably fast at >∼5 μm^2^/s (Fig. 4b), suggesting that a liquid-liked core remained even as the granule surface was covered by a thickened shell of amyloid aggregates.

## Conclusions

To sum, harnessing a set of multidimensional SRM tools, we unveiled nanoscale heterogeneity in FUS condensates, identifying hydrophobic amyloid aggregates at the condensate surface, showing their substantial suppression of local diffusivity, and elucidating their gradual expansion during aging.

For the LLPS of a single protein, one may intuitively assume each condensate microdroplet as a uniform phase. However, recent theoretical and experimental work points to the possible gradual transformation of initially homogenous FUS LLPS condensates to a liquid-core/gel-shell architecture through aging^19,25^. In this study, we uncovered nanoscale amyloid fibril aggregates at the condensate surface right after the condensates were formed, and showed that over aging, such nanoaggregates gradually expanded at the surface and increased their intra-condensate presence (model in Fig. 4d). These findings benefited from both the high spatial resolution and the multidimensional insights afforded by our approaches: Indeed, SMLM of Cy3B showed relatively uniform spatial distribution in the condensates, but SM*d*M of the same dye showed drastically impeded diffusion at the surface nanoaggregates, and SR-SMLM with Nile Red highlighted their hydrophobic nature and prompted us to employ CRANAD-2 SMLM to identify their amyloid state.

The low-complexity domains of FUS are known to drive both LLPS and fibril formation^11,12,53-58^. As FUS molecules aggregate into the amyloid state, their low-complexity domains are buried into a hydrophobic core, as detected by our Nile Red SR-SMLM. It is thus likely that the aggregated molecules no longer participate in the LLPS interactions and so are excluded from the condensate interior to appear on the surface. The observed gradual expansion of amyloid fibril nanoaggregates during aging, as well as their markedly enhanced formation upon repeated pipetting, suggest that while the LLPS condensates are initially fast formed under kinetic control, the fibril state is energetically favored as the system evolves toward equilibrium.

Together, our results unveiled unexpected structural arrangements and aging mechanisms for the single-component FUS condensates with exceptional spatial and functional insights. Generalizing the multidimensional SRM approaches demonstrated in this work to other LLPS systems to uncover related, or yet other unknown, behaviors at the nanoscale presents exciting opportunities.

## Acknowledgments

We acknowledge support by the National Institute of General Medical Sciences of the National Institutes of Health (DP2GM132681), the Packard Fellowships for Science and Engineering, and the Heising-Simons Faculty Fellows Award.

## Materials and Methods

### Plasmid constructs

FUS was PCR-amplified from pDEST8_FLAGHA_FUS_HIS (Addgene plasmid no. 26375; a gift from Thomas Tuschl)^59^. The baculovirus transfer vector pFastBac HT was PCR-amplified from pFastBac HT JS-Munc18b (Addgene plasmid no. 135554; a gift from Jingshi Shen)^60^. pFastBac HT 10xHis-MBP-TEV-FUS was constructed by the Gibson assembly (New England BioLabs E2611) of pFastBac HT, PCR-amplified 10xHis-MBP (a gift from Eunyong Park), and PCR-amplified FUS. The FUS(G156E) mutation was generated by changing the glycine codon (GGA) to a glutamic acid codon (GAA) using PCR between the BamHI and XbaI sites. All constructed plasmids were prepared from XL1-Blue cells using the QIAprep Spin Miniprep Kit (QIAGEN). Protein-coding sequences were verified by Sanger sequencing at UC-Berkeley DNA Sequencing Facility. Baculoviruses carrying the above constructs were generated using the Bac-to-Bac Baculovirus Expression System (Thermo Fisher) following the provider’s instructions.

### Protein expression

The above 10xHis-MBP-TEV-FUS and 10xHis-MBP-TEV-FUS(G156E) baculovirus constructs were expressed in 500 mL Sf9 insect cells. After 96 hr, pellet was collected by spinning the cells at 2,000 rpm for 5 min. From here, all the purification steps were done at 4°C until the protease cleavage step. The pellet was resuspended into 40 mL lysis buffer containing 50 mM Tris-HCl pH 7.4, 1 M KCl, 5% glycerol, and 10 mM imidazole. Protease inhibitors were added (Halt Protease Inhibitor Cocktail, Thermo Scientific 78429, excluding EDTA). The resuspended cells were lysed by sonication and clarified by centrifugation at 13,000 rpm for 30 min. The supernatant was collected and filtered with Corning syringe filters with a diameter of 28 mm and a pore size of 0.45 μm. The filtered supernatant was loaded onto 2 gravity columns each with 4 mL Ni-NTA agarose (Qiagen). The resin was pre-equilibrated with the lysis buffer. The protein was eluted with a buffer containing 50 mM Tris-HCl pH 7.4, 1 M KCl, 5% glycerol, and 250 mM imidazole. TEV protease (Sigma T4455) was added to the elute at a 1:50 ratio. The mixture was incubated at RT for 6 hr and then purified *via* gel filtration chromatography (ÄKTA go with Superdex-200 Increase 10/300 column, Cytiva) with the storage buffer containing 50 mM Tris-HCl pH 7.4, 500 mM KCl, 1 mM DTT and 5% glycerol. The peak fractions containing pure protein were concentrated by Amicon Ultra centrifugal filters (0.5 mL; 10 kDa MWCO) to ∼1 mg/mL, aliquoted into PCR tubes, flash-frozen by liquid nitrogen, and stored at −80°C.

### Glass surface preparation for microscopy

The protocol was adapted from Gidi *et al*^*61*^ and Nakashima *et al*^*62*^. #1.5 coverslips 12 mm in diameter and 24 × 50 mm in size were treated with a heated 3:1 H_2_SO_4_ (98%) and H_2_O_2_ (30%) mixture for 25 minutes, rinsed until neutral using Milli-Q water. The cleaned coverslips were sonicated in 0.5 M NaOH for 30 min, and rinsed until neutral using Milli-Q water. The coverslips were then sonicated in HPLC-grade acetone for 5 min, dried with N_2_, and then placed in a covered Petri dish in an oven at 60°C. Methoxy-PEG silane (MW 5000, PG1-SL-5k, Nanocs) was dissolved in dry DMSO at 30 mg mL^−1^ at 60°C. The solution was added to the top side of the coverslips and incubated at 60°C for 2 hr. Then, the coverslips were washed thoroughly with ethanol, MilliQ water (with 5 min sonication), and ethanol, dried with N_2_, and then kept in the oven.

### Preparation of condensate samples

An aliquot of the above FUS or FUS(G156E) stocks was thawed and spun at 13,000 rpm for 5 min. The supernatant was mixed at a 3:2 ratio with a crowding buffer [20% dextran (from *Leuconostoc* spp., Mr 450,000-650,000; Sigma 31392) and 1 mM DTT in 50 mM Tris-HCl pH 7.4]. Single or combined dyes were pre-dissolved in the crowding buffer, including 1 μM of Nile Red (415711000, Acros Organics), 10 μM of CRANAD-2 (4803, Tocris), and 5 nM of Cy3B hydrolyzed from Cy3B-NHS ester (PA63101, Cytiva). The final mixture thus contained ∼0.6 mg/mL FUS or FUS(G156E), 8% dextran, 300 mM KCl, 1 mM DTT, 3% glycerol in 50 mM Tris-HCl pH 7.4, and desired dyes. The sample was mixed thoroughly at room temperature on a bench rotor for 2 hr, and sealed between two functionalized coverslips as described above before being imaged. For aging over days, the sample was kept in the dark at room temperature without further agitations.

### Optical setup

SMLM, SR-SMLM, and SM*d*M were performed on a home-built setup based on a Nikon Ti-E inverted fluorescence microscope, as described previously^34,36^. Briefly, a 642-nm laser (Stradus 642, Vortran, 110 mW) and two 561-nm lasers (OBIS 561 LS, Coherent, 150 mW and 200 mW) were coupled to the back focal plane of an oil-immersion objective lens (Nikon CFI Plan Apochromat lambda 100x, NA 1.45). A translational stage shifted the laser to the objective edge to enter the sample slightly below the critical angle of total internal reflection. The sample was thus illuminated and imaged in the wide field a few micrometers away from the coverslip surface for a cross-section of the condensates.

### SR-SMLM with Nile Red

For SR-SMLM with Nile Red^34^, the sample was excited with the 561-nm laser at ∼3 kW/cm^2^. Wide-field emission was filtered with a long-pass filter (ET575lp, Chroma) and a short-pass filter (FF01-758/SP, Semrock), and cropped at the image plane to ∼4 mm in width. The cropped intermediate image was collimated by an achromatic lens before being split into two paths with a 50:50 beam splitter (BSW10, Thorlabs). One of the two split paths was passed through an equilateral prism (PS863, Thorlabs) for spectral dispersion, and both paths were imaged onto two different halves of an EM-CCD camera (iXon Ultra 897, Andor) that recorded continuously in the frame-transfer mode at 110 frames per second (fps). The stochastic encountering of individual Nile Red molecules with hydrophobic phases in the sample led to bursts of single-molecule fluorescence emission, for which the above system concurrently recorded the images and spectra of the same molecules in the wide field. ∼10^4^ frames were thus recorded, and the accumulated single-molecule images and spectra were processed into SR-SMLM images, in which color was used to present the mean wavelength of local single-molecule spectra^33,34^.

### SM*d*M with Cy3B

For SM*d*M with Cy3B^36,39^, a multifunction I/O board (PCI-6733, National Instruments) read the exposure timing of the EM-CCD camera, which recorded continuously at 110 fps, and accordingly modulated the 561-nm lasers to output tandem pulses across odd and even frames (Fig. 3a). Typical center-to-center separation between the paired pulses was Δ*t* = 4 ms, whereas the duration of each pulse was *τ* = 500 μs. The estimated peak and average power densities at the sample were ∼15 and ∼0.8 kW/cm^2^, respectively. Wide-field fluorescence emission was filtered by a long-pass filter (ET575lp, Chroma) and a band-pass filter (ET605/70m, Chroma). Single-molecule images due to Cy3B molecules stochastically diffusing into the focal plane were thus captured during the pulse illumination time *τ*, and their displacements were evaluated over the fixed time interval Δ*t* between the paired pulses. The above tandem excitation and recording scheme was repeated ∼10^4^ times, and the accumulated single-molecule displacements were spatially binned with a 120×120 nm^2^ grid. To generate super-resolved maps of the local diffusion coefficient *D*, the distribution of displacements in each spatial bin was separately fitted through maximum likelihood estimation (MLE) to a modified random-walk model with the probability distribution

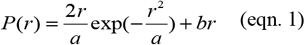

where *r* is the single-molecule displacement, *a* = 4*D*Δ*t*, and *b* is a background term to account for irrelevant molecules that randomly diffuse into the view, as rationalized and validated in ^36^. The fitted *D* values were color plotted on a continuous scale to generate spatial maps. To quantify the apparent two-component single-molecule displacements of Cy3B in the FUS-condensate interior, in this study we further introduce a new fitting model with the probability distribution

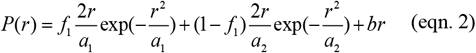

where *f*_1_ and *f*_2_ = (1− *f*_1_) are the fractions of the two diffusion components, and *a*_1_ = 4*D*_1_Δ*t* and *a*_2_ = 4*D*_2_Δ*t* account for the two diffusion coefficients *D*_1_ and *D*_2_.

### SMLM with CRANAD-2

For SMLM of CRANAD-2, the sample was continuously illuminated with the 642-nm laser at ∼4 kW/cm^2^. Wide-field emission was filtered by a long-pass filter (ET655lp, Chroma) and a band-pass filter (ET705/100m, Chroma), and recorded with the EM-CCD camera at 110 fps. Bursts of single-molecule fluorescence emission were achieved as CRANAD-2 molecules stochastically bound and unbound from amyloid fibrils^50^. The camera recorded ∼80,000 frames for each sample, and the localized single-molecule positions were assembled into SMLM images as described in ^63,64^. For two-color imaging, the sample was first imaged for CRANAD-2 with the 642-nm excitation with the above emission filters. The filters were then switched to the above respective sets for Nile Red SMLM or Cy3B SM*d*M under 561-nm excitation.

---

**Fig. S1.**
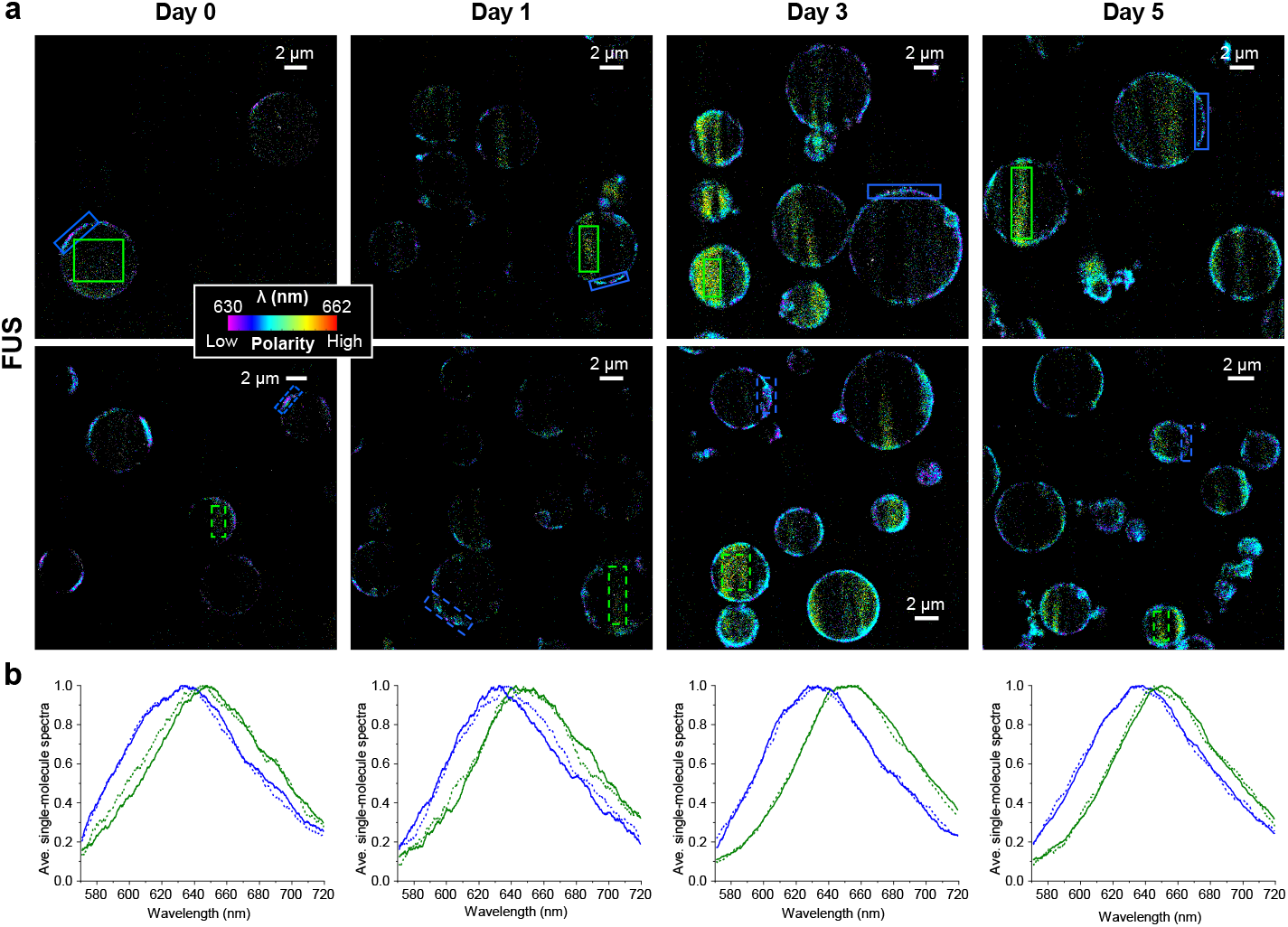
Additional Nile Red SR-SMLM data for condensates of wild-type FUS. **a**. Color-coded Nile Red SR-SMLM images of FUS condensates aged for different days. Two representative images are shown for each timepoint in addition to the images shown in Fig. 1b. Color presents the mean wavelength of local single-molecule spectra (color scale bar), which reflects the local chemical polarity. **b**. Averaged single-molecule spectra at the condensate surface hydrophobic domains (blue) and the condensate interior (green) at different timepoints, for the sold-line and dash-line boxed regions in (a). Note that the highly hydrated condensate interior results in both substantially redder (longer wavelength) and weaker Nile Red single-molecule emission. Consequently, fewer Nile Red molecules are detected in the condensate interior in the SR-SMLM data. However, as we illuminated the sample at a slanted angle close to the critical angle of total internal reflection (methods), scattered regions of enhanced illumination and hence single-molecule emission are noted, attributable to local lensing effects owing to the higher index of refraction of the surface aggregates unveiled in this study. Higher counts of single molecules are obtained for such regions.

**Fig. S2.**
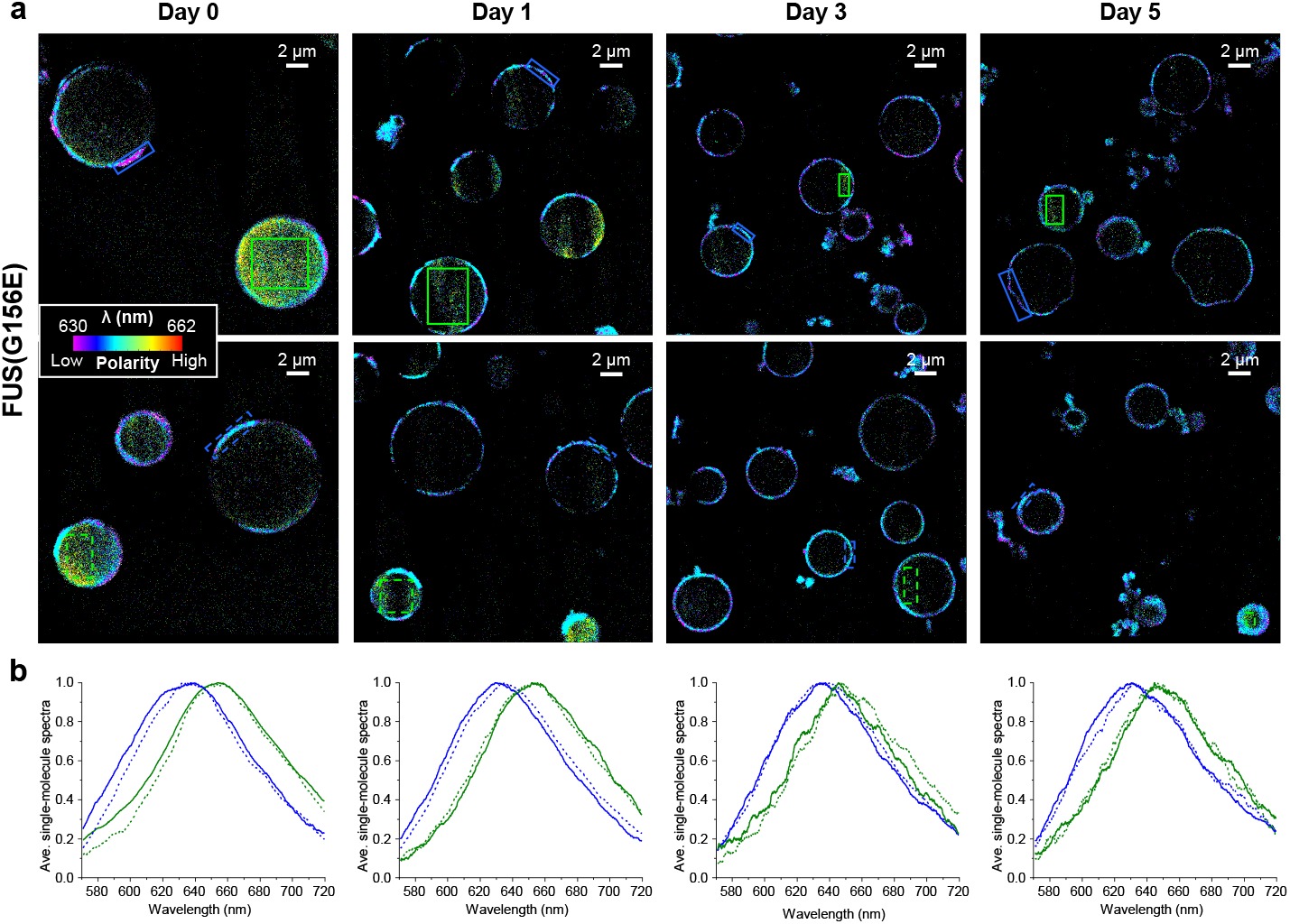
Additional Nile Red SR-SMLM data for FUS(G156E) condensates. Figure is arranged the same way as Fig. S1.

## Notes

### Competing Interest Statement

The authors have declared no competing interest.

